# IL-33 promotes transcriptional and metabolic adaptations of tissue-resident Th2 cells

**DOI:** 10.1101/2025.07.09.663905

**Authors:** Anna K. Kania, David E. Sanin, Xinyue Gu, Mia Gidley, Eryk Kokosinski, Allen Smith, Erika L. Pearce, Edward J. Pearce

**Affiliations:** Bloomberg Kimmel Institute of Cancer Immunotherapy, Department of Oncology, Johns Hopkins University School of Medicine, Baltimore, MD 21287, USA; USDA, Beltsville Human Nutrition Research Center and Diet, Genomics, and Immunology Laboratory, Beltsville, MD 20705, USA; Department of Molecular Microbiology and Immunology, Bloomberg School of Public Health, Johns Hopkins University, Baltimore, MD 21287, USA

## Abstract

The polarization of naïve CD4^+^ T cells into Th2 cells is initiated in lymphoid organs and completed as the cells become tissue resident, where they express ST2, the receptor for the alarmin IL-33, which may be a key signal for tissue integration. Cellular metabolic requirements associated with this transition remain poorly understood. To address this, we compared the response of lymphoid tissue (LT) Th2 cells from helminth parasite-infected mice to stimulation by IL-33 versus through the T cell receptor via anti-CD3/CD28. We found that IL-33, but not anti-CD3/CD28, induced the development of tissue-resident like Th2 cells expressing ST2. This was associated with IL-33 induced changes in arginine metabolism linked to mTORC1 activation and polyamine synthesis, which were required for the development of tissue-resident like Th2 cells. Futhermore, IL-33 induced transcriptional changes in genes involved in chemotaxis and cell adhesion that may be critical for tissue integration. Our findings provide insights into adaptations of Th2 cells responding to tissue-integration cues.

**Summary:** IL-33 promotes development of tissue-resident-like Th2 cells *in vitro* from lymphoid tissue Th2 cells. This requires arginine-dependent mTORC1 activation and polyamine synthesis, and is marked by transcription of genes associated with chemotaxis and cell adhesion linked to tissue integration.

## INTRODUCTION

Type 2 immunity is induced in response to helminth infections, where it can play a protective role, but also underpins allergic responses (1, 2). Type 2 immunity is largely controlled by cytokines released from T helper type 2 (Th2) cells and type 2 innate lymphoid cells (ILC2), both of which express the lineage-defining transcription factor GATA3 and secrete the canonical type 2 cytokines IL-4, IL-5, and IL-13 (3, 4). Naïve CD4^+^ T cell differentiation into Th2 cells is initiated in lymph nodes following TCR stimulation, and this commitment has been extensively modeled *in vitro* by activating naïve CD4^+^ T cells using anti-CD3/anti-CD28 antibodies, to engage the TCR and the costimulatory receptor, or antigen, in the presence of IL-4 and anti-IFNγ antibody. Under these conditions, the cells that develop express *Il4*, *Il5* and *Il13*. However, they do not express the hallmark gene of tissue resident Th2 cells, *IL1rl1*, which encodes the IL-33 receptor ST2, and do not secrete Amphiregulin (AREG), an EGFR1 ligand associated with tissue remodeling. This reflects the critical role for additional signals, provided by the tissue-derived alarmins IL-33, IL-25 and TSLP, in the terminal differentiation of Th2 cells as they become tissue-resident (4, 5). IL-33 is also implicated in the establishment of tissue resident regulatory T cells and CD8^+^ T cells and may represent a primary signal for tissue integration (6–8). Tissue-resident Th2 cells have been identified in the lamina propria, peritoneal cavity, and mesenteric adipose tissue (mAT) following infection with the parasitic helminth *Heligmosomoides polygyrus bakeri (Hpoly)* (9–11), as well as in inflamed lung and skin (12–14). However, due to their limited numbers, relatively little is known about the metabolic adaptation of these cells and the roles that distinct alarmins play in establishing their tissue residency.

Th2 cells play a critical role in the pathology of asthma, allergic rhinitis, and atopic dermatitis (15). In these cases, pathogenic Th2 cells are tissue-resident memory cells characterized by their ability to produce large amounts of IL-5 and IL-13. Similar to their intestinal counterparts, tissue-resident lung Th2 cells also express ST2 and can respond to alarmins secreted upon subsequent allergen exposure (16, 17). The persistence of these tissue-resident memory Th2 cells contributes to chronic inflammation and disease recurrence in allergic disorders, making them attractive therapeutic targets for conditions characterized by type 2 inflammation.

We found that stimulating lymphoid tissue-derived CD4^+^ T cells from *Hpoly*-infected mice (LT-Th2 cells) with IL-33 results in the expansion of a subset of cells expressing ST2, and the acquisition by these cells of new functions such as the increased expression of a panel of chemokines, chemokine receptors and adhesion molecules consistent with a requirement to integrate within tissues. These changes were not induced by stimulation through the TCR. We found IL-33-stimulated LT-Th2 cells to be highly metabolically active and dependent on arginine for proliferation and differentiation, reflecting critical roles for this amino acid in independently permitting mTORC1 activation and polyamine synthesis in these cells.

## RESULTS

### IL-33 stimulates LT-Th2 cells to express markers of tissue-resident Th2 cells

We infected wild-type mice with *Hpoly*, allowing the infection-driven initial step of Th2 cell differentiation to occur within lymphoid organs *in vivo*. At 2 wks post-infection, we isolated CD4^+^ T cells from secondary lymphoid organs (spleen and mesenteric lymph nodes) and cultured them *in vitro* with IL-2 (as a survival factor) alone, or with IL-33 or anti-CD3/anti-CD28 antibodies to provide alarmin-driven vs. TCR-driven activation signals. We included IL-4 and anti-IFNγ antibody in the anti-CD3/CD28 cultures to provide known Th2 cell promoting signals (Fig. S1A; hereafter referred to as anti-CD3/CD28). We detected IL-33 receptor ST2-positive GATA3^+^ cells after IL-33 stimulation (Fig. 1A), that were present at very low frequency immediately ex-vivo (Fig. S1B) and after anti-CD3/CD28 stimulation (Fig. 1A). IL-33 stimualted cells upregulated ST2, which was also expressed by *bona fide* tissue-resident Th2 cells isolated from the mAT of *Hpoly* infected mice (Fig. 1A). Adding IL-33 to the anti-CD3/CD28-stimulated cultures led to a slight increase in GATA3^+^ST2^+^ cells, but the frequencies and numbers of these cells were significantly lower than in the cultures stimulated only with IL-33 (Fig. S1C), suggesting that TCR signaling, IL4, or anti-IFNy (all present in the anti-CD3/CD28 cultures), might suppress ST2 expression, or signaling through ST2. Likewise, stimulating splenocytes from infected (but not naïve) mice with *Hpoly* antigen (HES) resulted in marked expansion of the GATA3^+^ population, but these cells were ST2^NEG^ (Fig. S1D, and data not shown), supporting the view that TCR stimulation expands cells that are GATA3^+^ but not expressing ST2.

**Figure 1.**
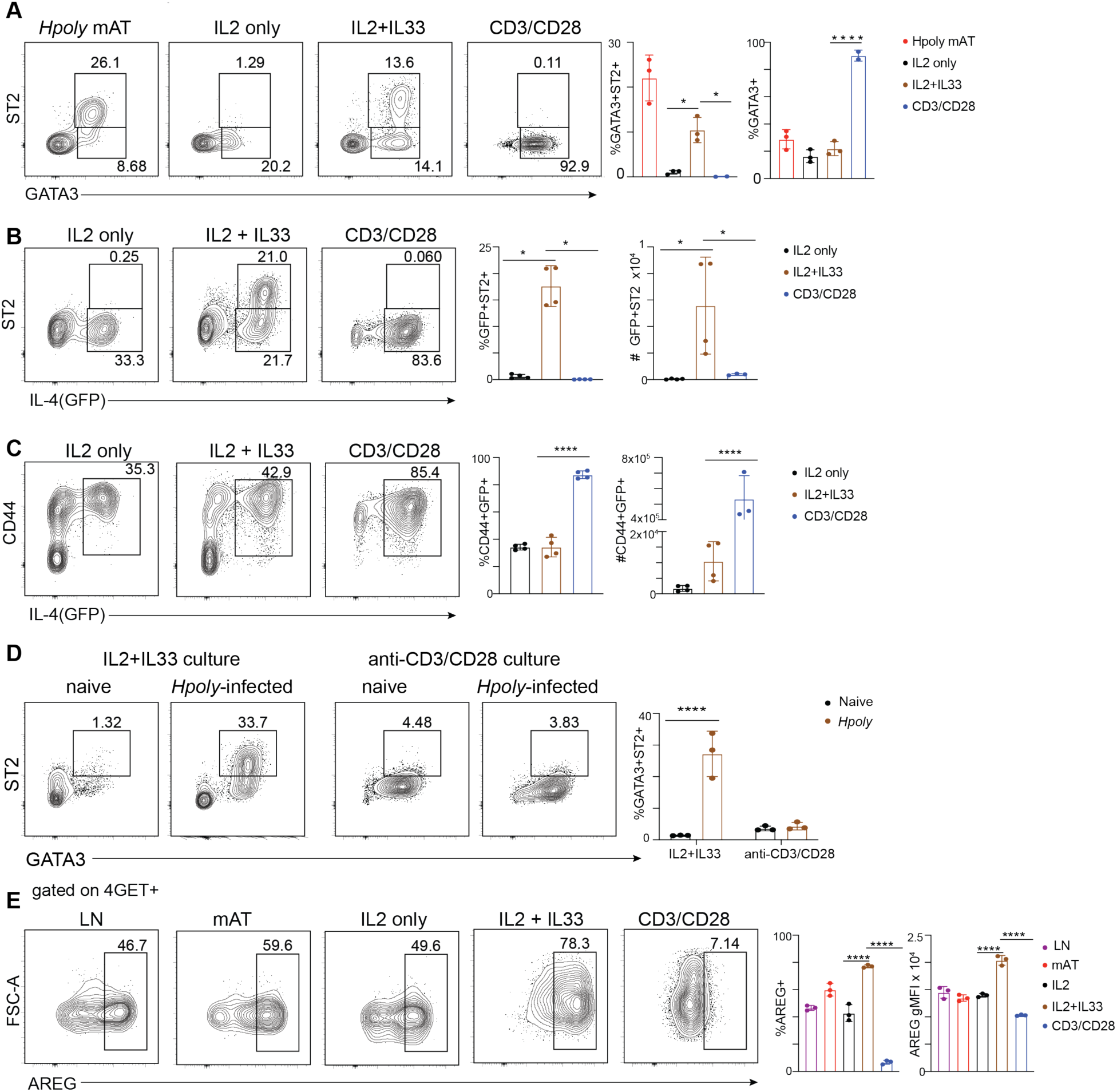
Generation of tissue-resident-like Th2 cells *in vitro*. (A) Representative flow plots and quantification of ST2+GATA3 + in live CD45+TCRβ+CD4+FOXP3-T cells from mesenteric adipose tissue of *Hpoly*-infected mice or after five days of *in vitro* culture of CD4 T cells isolated from Hpoly-infected mice. Flow plots and quantification of ST2+IL4+(GFP) (B) or CD44+IL4+(GFP+) (C) cells cultured as above. (D) CD4 T cells from naïve or *Hpoly*-infected mice were cultured as in (A) and IL-33R+GATA3+ Th2 cells were quantified. (E) Representative flow plots and quantification of AREG+ cells within the IL4+(GFP+) population after 5hr restimulation with PMA/Iono in the presence of Brefeldin. The data (A-E) are representative of at least two independent experiments. Symbols in the quantified data represent independent biological replicates. Data were analyzed by one-way ANOVA with Tukey’s post hoc test (A-C, E) or two-way ANOVA with Sidak post hoc test (D).. * p ≤ 0.05, ** p ≤ 0.01, *** p ≤ 0.001, *** p ≤ 0.0001.

We next assessed the effects of IL-33 vs. anti-CD3/CD28 on CD4^+^ T cells from secondary lymphoid organs of *Hpoly* infected 4get mice, in which *Il4* expression is reported by GFP production (18) (again using the protocol in Fig. S1A). This approach allowed us to focus on LT-Th2 cells in which *Il4* had been expressed. Directly after isolation, approximately 15% of the CD4^+^ T cells from *Hpoly*-infected mice expressed GFP (and therefore were IL-4^+^), although very few cells expressed ST2 (Fig. S1E). After five days of culture, we found that ST2 expression was apparent only when cells were stimulated with IL-33, and that only GFP^+^ cells expressed ST2 under these conditions (Fig. 1B). This was notable because anti-CD3/CD28 was nevertheless a potent stimulus for the expansion of GATA3^+^ (Fig. 1A) and GFP^+^ populations (Fig. 1C). The ability of IL-33 to induce the development of GATA3^+^ST2^+^ cells was dependent on CD4^+^ T cells having been Th2-primed *in vivo*, since very few GATA3^+^ST2^+^ cells emerged when CD4^+^ T cells from naïve mice were stimulated with IL-33 (Fig. 1D). Moreover, CD4^+^ T cells from naïve as well as infected mice upregulated GATA3 following stimulation with anti-CD3/CD28 under Th2 conditions, but neither population expressed ST2 (Fig. 1D). These data indicate that there is a critical in vivo event in responsive secondary lymphoid organs that allows the development of “LT-Th2” cells that are ready to respond to IL-33. However, this event is not recapitulated *in vitro* by stimulating naïve T cells with anti-CD3/CD28 under Th2 conditions. To gain a better understanding of the kinetics of the IL-33 driven response, we followed the development of GATA3^+^ST2^+^ cells over time. Again, CD4^+^ T cells from naïve mice failed to upregulate ST2 in response to IL-33. On the other hand, the frequencies and numbers of GATA3^+^ ST2^+^ Th2 cells progressively increased over time when CD4^+^ T cells from the secondary lymphoid organs of *Hpoly* infected mice were stimulated with IL-33 (Fig. S1F).

To assess the functionality of IL-33-stimulated Th2 cells, we stimulated cells with PMA/Iono and measured AREG using intracellular flow cytometry. AREG is a major product of tissue-resident Th2 cells (9, 19, 20). Consistent with this, ∼60% of PMA/Iono-stimulated IL4^+^(GFP^+^) Th2 cells recovered from the mAT of *Hpoly* infected 4get mice produced AREG (Fig. 1E). We found that after 5 days in culture, ∼50% IL4^+^(GFP^+^) cells expressed AREG in response to IL-2, whereas ∼80% expressed this cytokine when the cells were additionally stimulated with IL-33 (Fig. 1E). Moreover, the MFI of staining of AREG was significantly increased after stimulation with IL-33 (Fig. 1E). Cells stimulated with anti-CD3/CD28 did not make AREG under these conditions (Fig. 1E). It was noticeable in these assays that both IL-33 and anti-CD3/CD28 stimulated increases in forward-scatter/cell size (Fig. 1E), indicative of increased anabolic metabolism.

Combinatorial exposure to IL-33, IL-25 and TSLP has been shown to be critical for the terminal differentiation of Th2 cells (4, 5). To assess whether the observed effects of IL-33 are shared by TSLP and IL-25, we stimulated CD4^+^ T cells from secondary lymphoid organs of *Hpoly* infected mice with IL-2 alone or with IL-33, TSLP or IL-25, or combinations of these alarmins, and assessed GATA3 and ST2 expression as well as AREG secretion. We found that the percentages of cells expressing GATA3 and ST2 were increased by IL-25, although not to the same extent as with IL-33, and this did not result in increased numbers of GATA3^+^ST2^+^ cells (Fig. S1G). TSLP did not promote the development of GATA3^+^ST2^+^ cells, and neither IL-25 nor TSLP alone synergized with IL-33 to promote the development of GATA3^+^ST2^+^ cells (Fig. S1G). Nevertheless, the combination of all three cytokines promoted the expansion of the GATA3^+^ST2^+^ population compared to IL-33 alone (Fig. S1G). Furthermore, while only IL-33 alone stimulated the production of AREG, the combination of all three alarmins did result in an increase in AREG production (Fig. S1H). The data from this analysis also highlights that LT-Th2 cells stimulated with IL2 alone are incapable of secreting AREG in response to IL-33 (Fig. S1H), consistent with the fact that IL-2 alone cannot induce high expression of ST2 (Fig. 1A,B), and therefore sensitivity to IL-33. This contrasts with the fact that IL-2 stimulated LT-Th2 cells do have the capacity to make some AREG when stimulated in an IL-33/ST2-independent manner with PMA/Ionomycin (showin in Fig. 1E).

We hypothesized that the increase in the frequency of GATA3^+^ST2^+^ cells following stimulation with IL-33 was the result of an expansion of the small population of weakly ST2-expressing LT-Th2 cells residing in the secondary lymphoid organs of infected mice (Fig. S1B). To test this, we FACS-isolated CD44^+^IL-4^+^(GFP^+^)CD4^+^ LT-Th2 cells as well as CD44^+^IL-4^NEG^(GFP^NEG^) CD4^+^ activated non-Th2 cells, and CD44^NEG^IL-4^NEG(^(GFP^NEG^) CD4^+^ naïve T cells from secondary lymphoid organs of *Hpoly*-infected mice. Immediately *ex-vivo*, the sorted LT-Th2 cells showed low expression of ST2 (Fig. S2A, B). After 5 days of stimulation with IL-2 and IL-33, ∼77% of the cells were strongly ST2^+^ (Fig. S2A, B) and there was no decline in the percentage of CD44^+^IL-4^+^(GFP^+^) cells over this time (Fig. S2A). A subset of the cells that were sorted as CD44^+^IL-4^NEG^(GFP^NEG^) at the beginning of the culture upregulated GFP and ST2 in response to IL-33, probably reflecting the expansion of the small population of GFP^LOW^ST2^LOW^ cells that contaminated this sort (Fig. S2A, B). CD44^NEG^IL-4^NEG(^(GFP^NEG^) cells did not respond to IL-33 in this assay. (Fig. S2A, B). These data indicate that IL-33 functions largely by driving the expansion of a small population of ST2-expressing cells within the LT-Th2 cell pool.

### The IL-33-induced development of tissue-resident-like Th2 cells is mTORC1 dependent

To explore the metabolic effects of IL-33, we performed RNA-seq on IL4^+^ (GFP^+^) LT-Th2 cells immediately after isolation from *Hpoly*-infected mice and after five days in culture with IL-2 alone or together with IL-33, or anti-CD3/CD28 (Fig. S1A). Principal component analysis revealed extensive, distinct changes in gene expression in response to IL-33 vs anti-CD3/CD28 (Fig. 2A), and hierarchical clustering of differentially expressed genes confirmed that IL-33-stimulated cells exhibited a unique transcriptional signature (Fig. 2B).

**Figure 2.**
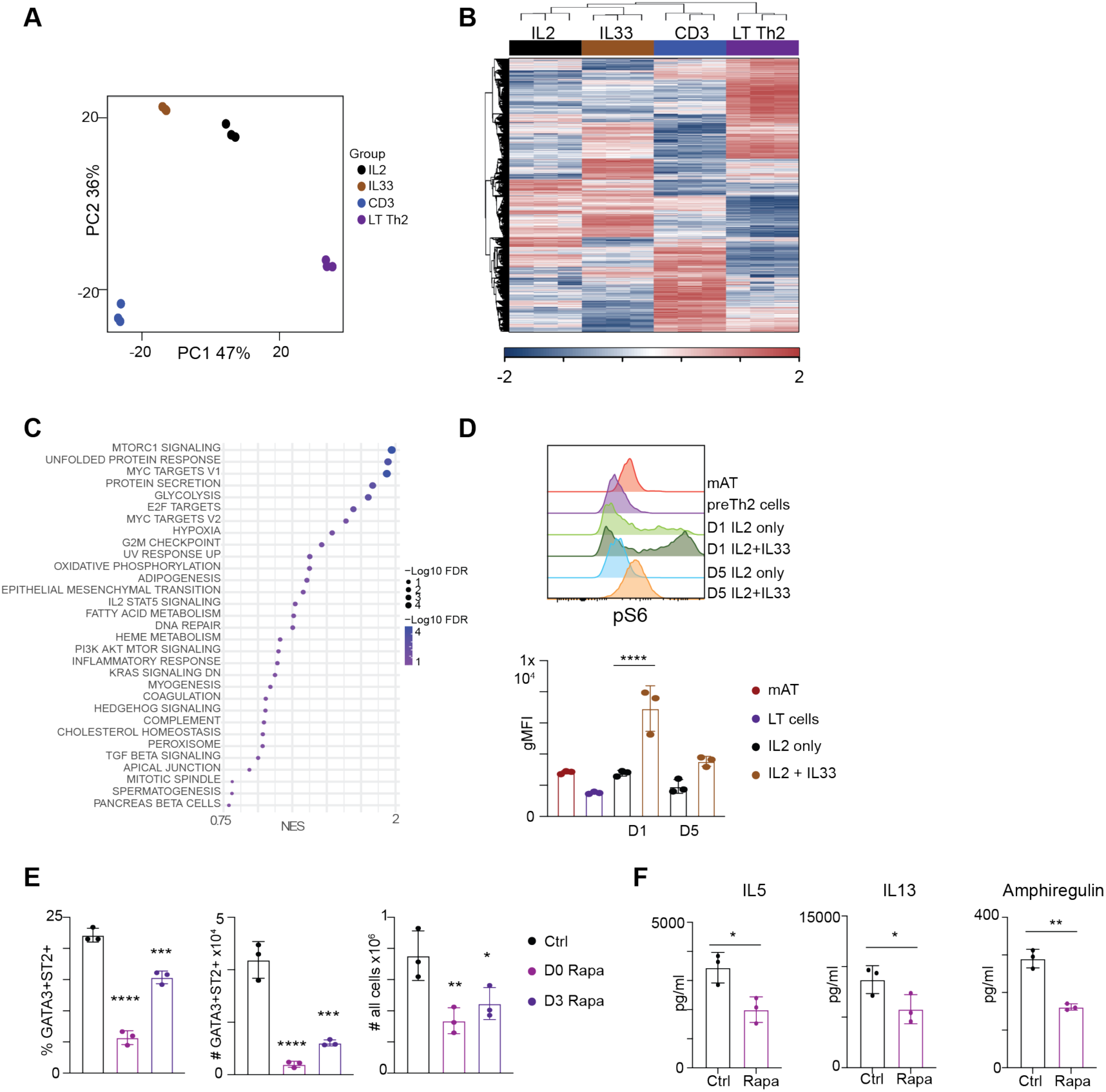
Th2 cells require mTORC for optimal differentiation. RNAseq was performed on CD44+IL4+(GFP+) Th2 cells isolated from spleens and lymph nodes of *Hpoly*-infected mice or after five-day culture under indicated conditions. (A) Principal component analysis of RNAseq data. (B) Heatmap of hierarchical clustering of differentially expressed genes. (C) Top Hallmark pathways from GSEA for IL-33 v IL2 comparison. (D) Flow plots and quantification of pS6 expression in CD44+IL4(GFP+) Th2 cells. (E) CD4 T cells were isolated from *Hpoly*-infected mice and cultured with IL2 and IL-33 in the presence of rapamycin. Frequencies and numbers of GATA3+ST2+ Th2 cells were quantified. (F) Levels of indicated cytokines after overnight culture in fresh media containing rapamycin. The data (D, E, F) are representative of at least two independent experiments. Symbols in the quantified data represent independent biological replicates. Data were analyzed by one-way ANOVA with Tukey’s post hoc test (D, E) or paired T-test (F). * p ≤ 0.05, ** p ≤ 0.01, *** p ≤ 0.001.

To gain further insight into pathways regulated by IL-33, we performed Gene Set Enrichment Analysis (GSEA) using HALLMARK gene sets. This revealed a highly significant enrichment of genes associated with mTORC1 signaling in IL-33-stimulated cells (Fig. 2C, S3A). Consistent with this, IL-33 stimulated cells showed positive staining for S6 phosphorylation (phospho-S6), a downstream consequence of mTORC1 activation (Fig. 2D). Futhermore, phospho-S6 staining in cells stimulated with IL-33 for 5 days resembled the level of *ex-vivo* phospho-S6 staining in IL-4^+^(GFP^+^) mAT tissue-resident Th2 cells from *Hpoly*-infected 4get mice (Fig. 2D). To further probe the role of mTORC1 in IL-33-driven differentiation of tissue-resident-like Th2 cells, we stimulated LT-Th2 cells isolated from the secondary lymphoid organs of *Hpoly*-infected mice with IL-33 in the presence or absence of 20 nM Rapamycin, which at this concentration selectively inhibits mTORC1 while sparing mTORC2. Rapamycin was added either at the initiation of culture, or at day 3, and cells were analyzed at day 5. Inhibiting mTORC1 significantly impaired the emergence of GATA3^+^ST2^+^ cells (Fig. 2E). A similar effect was observed when anti-CD3/CD28-stimulated cells were treated with rapamycin, where a significant reduction in GATA3^+^ cells was observed (Fig. S3B). These results are in line with the known role of mTORC1 in promoting cell cycle entry and proliferation (21, 22). Rapamycin also inhibited cytokine production by LT-Th2 cells that had been stimulated in vitro with IL-33 ((Fig. 2F) or anti-CD3/CD28 (Fig. S3C) for 5 days and then cultured overnight in fresh media in the presence of absence of the drug. Taken together, the data supports previous findings on the critical role of mTORC1 for cell proliferation and Th2 cell differentiation and function (22).

Along with the mTORC1 pathway, GSEA analysis also revealed significant upregulation of genes associated with oxidative phosphorylation and glycolysis, findings that are consistent with the reported metabolic changes induced by IL-33 in human ILC2s (23). To directly assess metabolic activity, we used SCENITH (24) on LT-CD4^+^ T cells from *Hpoly*-infected mice after stimulation for five days with IL-2 alone or with IL-33 or anti-CD3/CD28 (as in Fig. S1A). With SCENITH, protein translation, measured using puromycin incorporation, is used as a proxy for ATP synthesis and measured after treatment with oligomycin (to inhibit ATP production by mitochondria), or 2-deoxyglucose (2DG) (to inhibit glycolysis). We found that regardless of stimulation, GATA3^+^ cells had high glucose dependence but low mitochondrial dependence, suggesting that they are highly glycolytic (Fig. S3D). Furthermore, all cells showed comparable capacity to perform glycolysis and to oxidize fatty acids (FAO) or amino acids (AAO). To directly assess mitochondria, we measured mitochondrial mass and inner membrane potential after five days of culture. Since Mitotracker Green, which measures mass, is not compatible with cell fixation and permeabilization required for intracellular GATA-3 staining, and ST2 is not expressed by anti-CD3/CD28 stimulated cells, we gated on TSLPR^+^ cells to identify Th2 cells (Fig. S3E). We found that the frequency of cells with high mitochondrial mass was increased in TSLPR^+^ Th2 cells after IL-33 and anti-CD3/CD28 stimulation and that mitochondria with the greatest mass had the highest membrane potential (measured by Mitotracker DeepRed) (Fig. S3F). Indeed, the mitochondrial membrane potential of metabolically active IL-33-stimulated cells was slightly higher than in the anti-CD3/CD28 stimulated cells (Fig. S3F). Together, these results indicate that LT-Th2 cells cultured in IL-2 are metabolically active, and that stimuli that drive proliferation and cytokine production further potentiate metabolic activity.

### Arginine is necessary for Th2 cell differentiation

We used mass spectrometry to assess IL-33-induced metabolic changes in greater detail. For these experiments, we sorted IL-4^+^(GFP^+^) CD4^+^ LT-Th2 cells from infected 4get mice, as well as CD44^NEG^GFP^NEG^CD4^+^ T cells (naïve cells) from the same animals. In addition, we sorted IL-4^+^(GFP^+^) cells after five days of culture using the protocol shown in Fig. S1A. Metabolites in extracts of these sorted cell populations were analyzed by flow-injection mass spectrometry. Hierarchical clustering of the top 100 metabolites revealed metabolites that were enriched in the IL-33-stimulated cells, namely amino acids and metabolites involved in regulating oxidative stress (Fig. 3A,Table 1). To validate these findings, we performed LC/MS metabolomics analysis on naïve CD4^+^ T cells as well as IL-33-stimulated and anti-CD3/CD28-stimulated CD4^+^CD44^+^ cells. We examine the abundance of arginine, iso(leucine), and glutamine, which are known to promote mTORC1 activation (25) as well as proline, which can be used by cells to generate arginine. We found arginine was enriched in both activated cell groups compared to naïve cells (Fig. 3B). There were no significant differences in the abundance of proline or iso(leucine) but glutamine was significantly lower in IL-33-stimulated cells (Fig. 3B).

**Figure 3.**
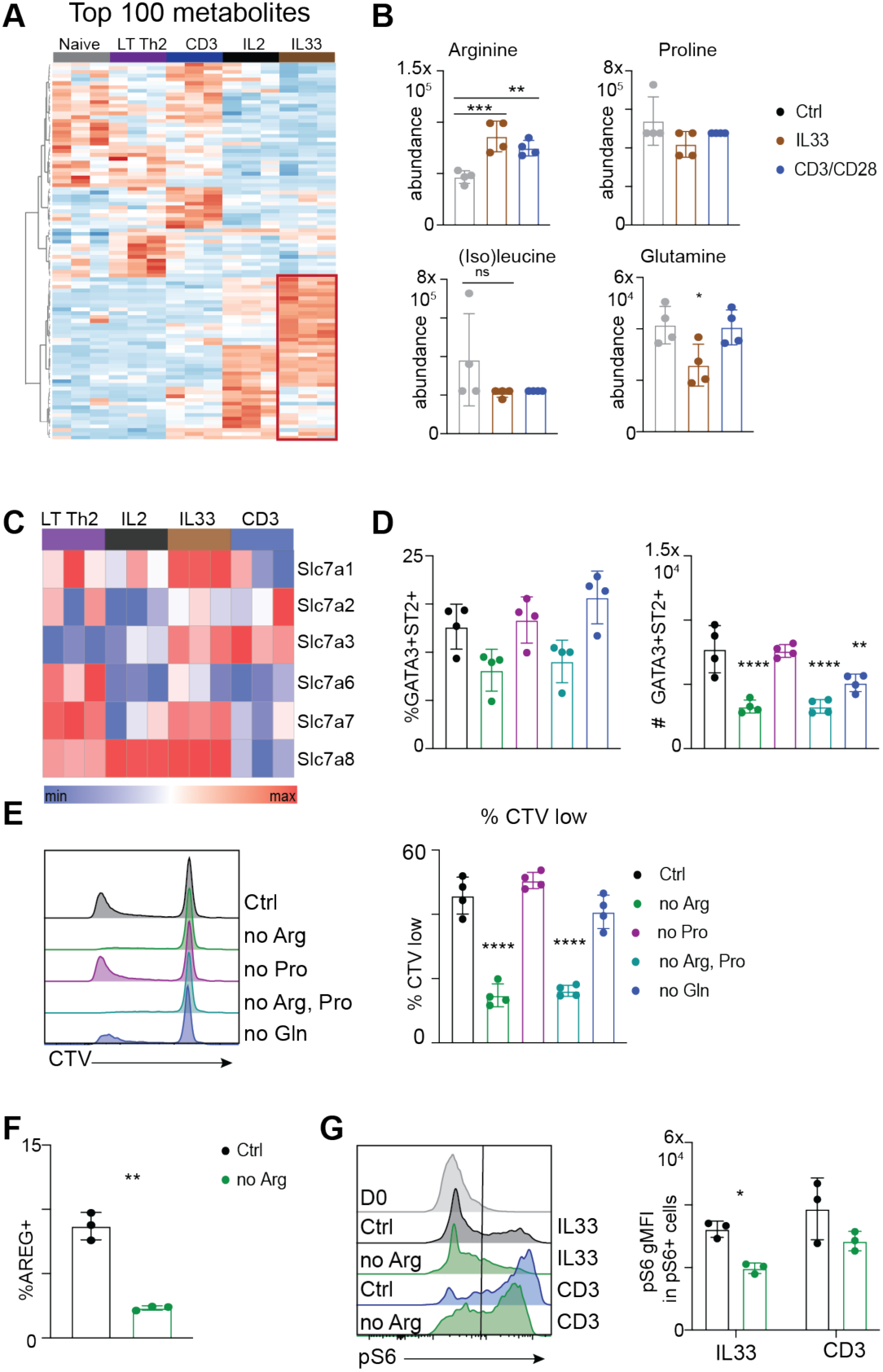
Arginine is required for Th2 cell differentiation. (A) Heatmap representing the abundance of the top 100 detected metabolites. (B) CD4 T cells were isolated from *Hpoly*-infected mice and cultured with IL2 and IL-33 for five days. The intracellular abundance of indicated amino acids in naïve and CD4+CD44+ activated cells was determined via LC/MS. (C) Heatmap depicting the expression of arginine transporters genes related in RNAseq data from Fig. 2. (D) CD4 T cells were isolated from *Hpoly*-infected mice and cultured with IL2 and IL-33 for five days in amino acid drop-out media. Frequencies and numbers of GATA3+ST2+ Th2 cells were quantified. (E) Representative histograms and quantification of CTV dilution of CD4 T cells cultured as in D. (F) After five days of culture with IL2 and IL-33 in control or arginine-depleted media, cells were restimulated with PMA/Iono and amphiregulin was quantified. (G) Representative histograms and quantification of ps6 levels in pS6-positive cells at day 1 of culture. The data (D-G) are representative of at least two independent experiments. Data were analyzed by one-way ANOVA with Tukey’s post hoc test (D, E), paired T-test (F), or two-way ANOVA with Sidak post hoc test. * p ≤ 0.05, ** p ≤ 0.01, *** p ≤ 0.001.

**Table 1.**
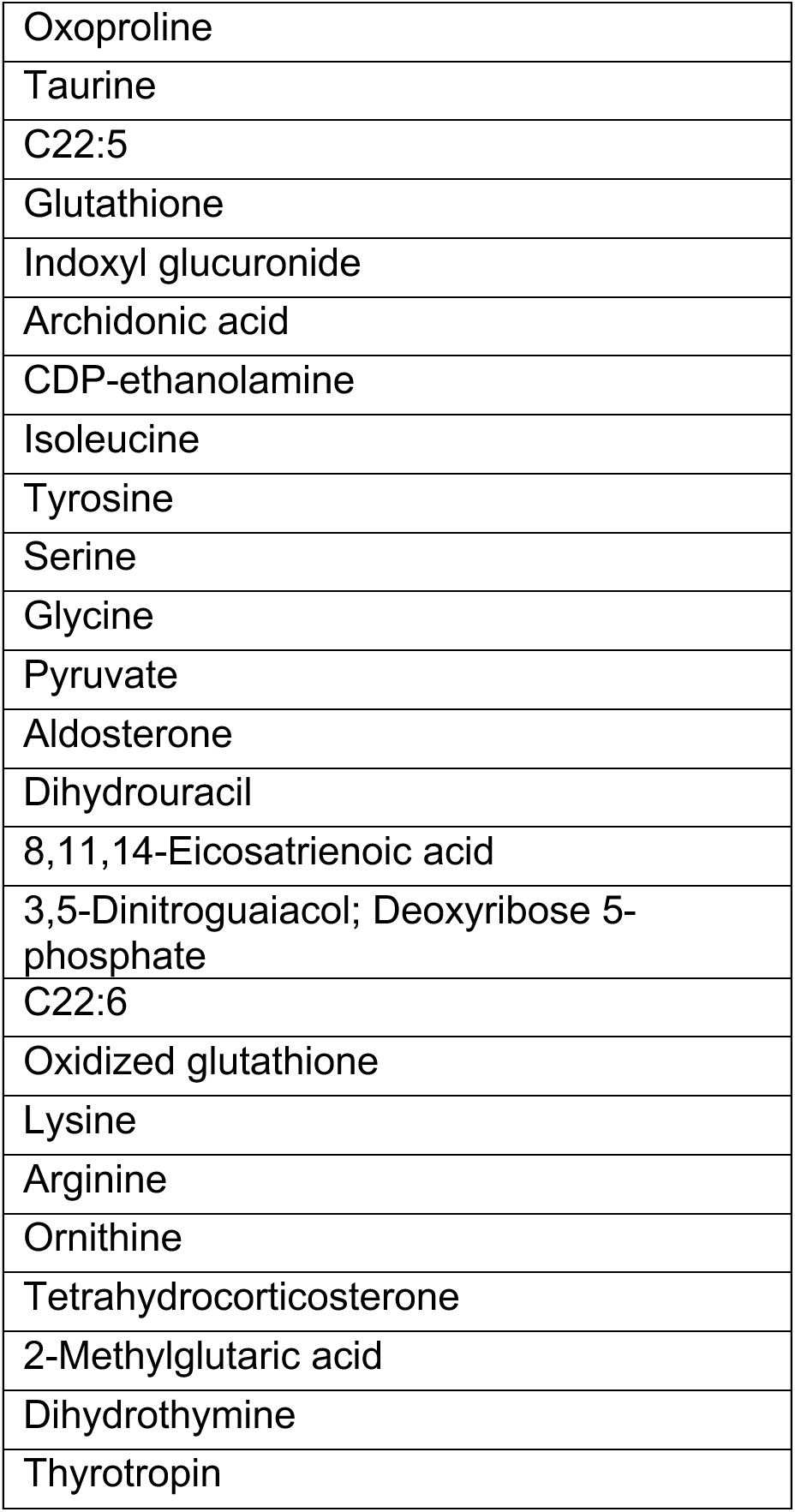
Metabolites upregulated in IL-33-stimulated Th2 cells red box in figure 3A.

It was of interest to us that arginine was amongst the amino acids enriched in IL-33-stimulated cells, since this is one of a select group of amino acids known to promote mTORC1 activation (26, 27). We reasoned that IL-33 may drive increased arginine uptake to support mTORC1 activation. Consistent with this, we found that the expression of genes encoding arginine transporters (28, 29) was upregulated in IL-33 stimulated cells (Fig. 3C). Of these, *Slc7a1* and *Slc7a8* were the most strongly expressed after IL-33 stimulation (Fig. 3C). To directly assess the role of arginine in terminal Th2 cell differentiation, we cultured LT-Th2 cells with IL-2 plus IL-33 in media lacking this amino acid, or proline or glutamine. Arginine restriction, but not restriction of either proline or glutamine, inhibited IL-33-driven GATA3^+^ST2^+^ Th2 cell differentiation (Fig. 3D), and cellular proliferation, as measured by the dilution of CTV (Fig. 3E). Despite the fact that proline has the potential to be used to generate arginine, depleting both amino acids did not have a synergistic effect (Fig. 3D, E). In keeping with its effect on the IL-33-driven emergence of ST2^+^ Th2 cells, arginine restriction additionally blunted the ability of IL-33 to promote AREG production (Fig. 3F). Glutamine depletion had no measureable effect on IL-33-driven proliferation or terminal differentiation (Fig. 3D, E). Finally, we assessed whether arginine restriction impaired mTORC1 activation. We found that withdrawal of arginine significantly reduced phospho-S6 in LT-Th2 cells stimulated with IL-33, but had a lesser (not statistically significant) effect on phospho-S6 in anti-CD3/CD28 stimulated cells (Fig. 3G), supporting the view that arginine uptake is particularly important for mTORC1 activation related to terminal Th2 cell differentiation.

### Arginine-fueled polyamine synthesis supports tissue-resident-likeTh2 cell differentiation

In addition to playing a role in mTORC1 activation, arginine is a substrate for polyamine synthesis. Polyamines are small cationic metabolites involved in cell replication, transcription, translation, and post-translational protein modifications (30, 31) and were recently shown to play a critical role in maintaining the epigenome to enforce Th cell subset fidelity (32, 33). We found that IL-33 induced the expression of genes encoding enzymes in polyamine metabolic pathways, including *Odc1* (Ornithine Decarboxylase), which converts ornithine to putrescine, *Dhps* (Deoxyhypusine Synthase), which facilitates eIF5A hypusination, and to a lesser extent *Arg1* (encodes Arginase 1), which converts arginine to ornithine (Fig. 4A, B). To assess whether these genes are also expressed in tissue-resident Th2 cells *in vivo*, we analyzed T cells within a scRNAseq data set of cells in the mAT of *Hpoly*-infected mice, i.e. tissue-resident Th2 cells (Fig. S4A) (9). We found that *Odc1*, *Srm*, *Dhps* and *Dohh* were also expressed within the GATA3^+^ST2^+^ cluster of cells in this dataset (Figs. 4C, S4A). To examine the role of polyamine metabolism in IL-33 induced terminal Th2 cell differentiation, we cultured LT-Th2 cells with IL-33 plus IL-2 or anti-CD3/CD28 containing ^13^C-arginine for the duration of the culture, and used LC/MS to assess the contribution of ^13^C to cellular arginine, putrescine, spermidine and spermine. We found that all intracellular arginine was ^13^C-labelled, and further detected significant labeling of three downstream polyamine metabolites (Fig. 4D), indicating that IL-33 stimulated cells do take up and metabolize environmental arginine for polyamine synthesis.

**Figure 4.**
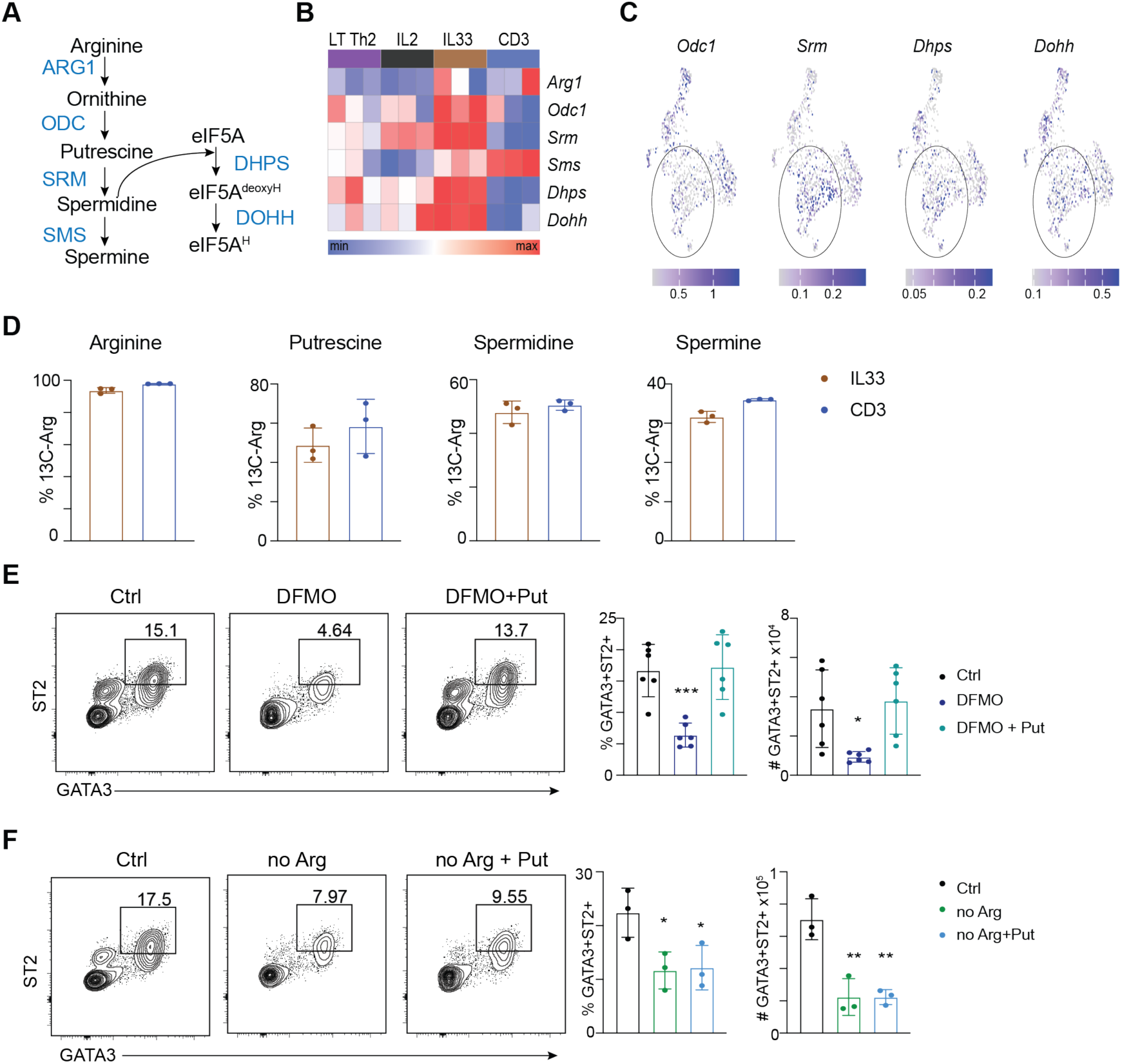
Th2 cells utilize arginine for polyamine synthesis. (A) Schematic of polyamine synthesis pathways. (B) Heatmap representing the expression of genes in the polyamine pathways in RNAseq data from fig 2. (C) Expression of selected genes from (B) in scRNAseq dataset of T cells isolated from mAT of *Hpoly*-infected mice. (D) CD4 T cells from Hpoly-infected mic were cultured with IL2 and IL-33 or CD3/CD28 in media containing 13C-Arg. Fractional contribution of labeled arginine to the indicated metabolites. CD4 T cells from *Hpoly*-infected mice were cultured with IL2+IL-33 for five days in the presence of DFMO (E) or in arginine-depleted media (F). The data (D-F) are representative of at least two independent experiments. Symbols in the quantified data represent independent biological replicates. Data were analyzed by one-way ANOVA with Tukey’s post hoc test (E-F). * p ≤ 0.05, ** p ≤ 0.01, *** p ≤ 0.001.

Similar results were observed for anti-CD3/CD28 stimulated cells (Fig. 4D). These results suggest that arginine uptake, rather than synthesis, is likely to be critical for tissue-resident Th2 cell differentiation. Next, we assessed whether inhibiting ODC would mimic arginine-restriction. Indeed, difluoromethylornithine (DFMO), an irreversible inhibitor of ODC, significantly impaired IL-33-driven tissue-resident-like GATA3^+^ST2^+^ Th2 cell development (Fig. 4E). Importantly, this effect was significantly ameliorated by supplementation of putrescine (Fig. 4E), which is the downstream product of the ODC-catalyzed reaction in the polyamine metabolism pathway (Fig. 4A). DFMO treatment also impaired the expansion of GATA3^+^ cells by of anti-CD3/CD28 (Fig. S4B). Of note, putrescine could not rescue impaired mature Th2 cell differentiation resulting from arginine restriction (Fig.4F, S4C). We speculate that this is due to the fact that putrescine cannot replace arginine-driven mTORC1 activation. Taken together, our data indicate that arginine supports IL-33-driven Th2 cell terminal differentiation through two distinct processes, permitting mTORC1 activation and in fueling polyamine metabolism.

### IL-33 promotes the expression of genes that encode proteins critical for cell movement and positioning within tissues

To identify IL-33-target genes in Th2 cells, we generated a list of genes (465 in total) that were upregulated in LT-Th2 cells following stimulation with IL-33 plus IL-2 versus IL-2 alone or versus anti-CD3/CD28 plus IL-2 (Fig. 5A). Gene Ontology analysis of this set of genes revealed significant associations with pathways such as inflammatory response and immune response (Fig. 5B), but also chemotaxis, cell adhesion, and positive regulation of cell migration (Fig. 5B), consistent with a role for IL-33 in promoting tissue-integration. Genes contributing to this association are shown in a heatmap in Fig. 5C. Furthermore, we found that *Ccl17, Ccr1, Ccr8,* and *Icam1* were also expressed in mAT-resident Th2 from *Hpoly* infected mice (Fig. 5D)(9). To validate these findings, we assessed the protein levels of CCR8 in mAT Th2 cells as well as IL-33 stimulated tissue resident like Th2 cells generated in vitro using our protocol. CCR8 is a chemokine receptor for CCL1 and CCL8, which promote cell migration to inflammatory sites, and was previously shown to be critical for Th2 cells in allergen-inflamed skin (34–36). Consistent with the transcriptional data, IL-33-stimulated cells expressed higher levels of CCR8, albeit not to the same extent as Th2 cells found in the mAT (Fig. 5E).

**Figure 5.**
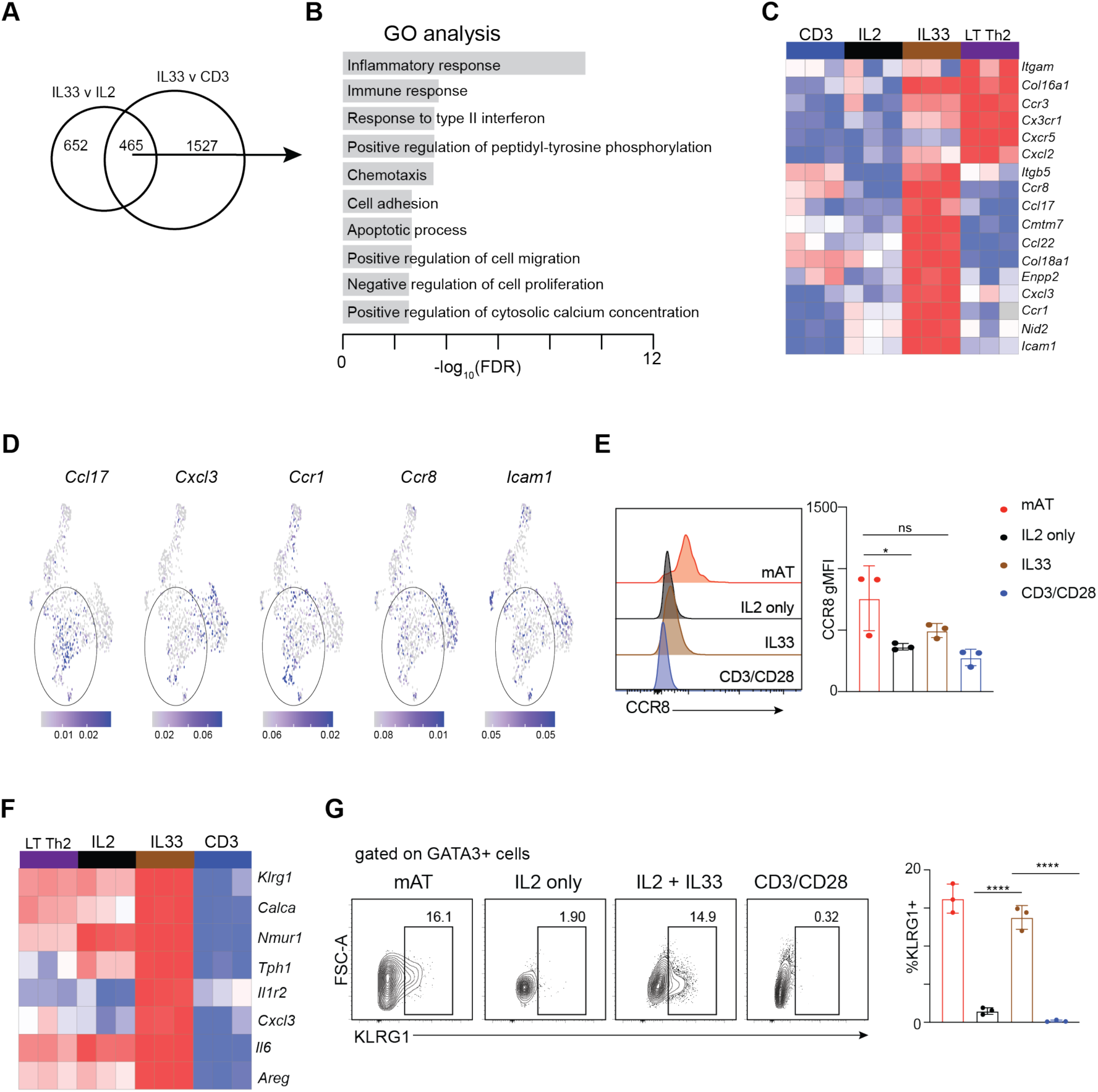
IL-33 induces expression of tissue-residency genes. (A) Venn diagram depicting the overlap of DEG between the two indicated comparisons. (B) Top Gene Ontology pathways enriched in the common 465 DEG from (A). (C) Heatmap showing the expression of selected genes in the chemotaxis and cell adhesion pathways from (B). (D) Expression of selected genes from (C) in scRNAseq dataset of T cells isolated from mAT of *Hpoly*-infected mice. (E) Representative histograms and quantification of CCR8 expression. (F) Heatmap showing the expression of known ILC2-related genes. (G) Representative histograms and quantification of KLRG1 expression. The data (E,G) are representative of at least two independent experiments. Symbols in the quantified data represent independent biological replicates. Data were analyzed by one-way ANOVA with Tukey’s post hoc test (E-F). * p ≤ 0.05, ** p ≤ 0.01, *** p ≤ 0.001.

Early work identified transcriptional overlap between ILC2s and tissue resident Th2 cells (5) and our data suggest that responsiveness to IL-33 is at least in part responsible for this, as we found increased expression of genes typically associated with ILC2s, including *Calca, Nmur1 and Klrg1, as well as Il1r2*, after LT-Th2 cells were stimulated with IL-33 (Fig. 5F). We confirmed increased KLRG1 expression in a subset of GATA3^+^ Th2 cells from the mAT of *Hpoly* infected mice, and from IL-33 stimulated LT-Th2 cell cultures (Fig. 5G). This was not observed when LT-Th2 cells were stimulated with IL-2 alone or with anti-CD3/CD28. Together, these data support the view that IL-33 promotes the expression of the transcriptional program required for tissue residency, and the expression of genes that are also expressed strongly in ILC2.

## DISCUSSION

We find that IL-33 is able to stimulate the development of lymphoid tissue Th2 cells into ST2^+^ AREG^+^ tissue resident-like Th2 cells in vitro, an effect that is not recapitulated by stimulation through the TCR. Despite the recognized importance of IL-25 and TSLP as type 2 immune alarmins, we found that IL-33 alone is able to directly induce the development of LT-Th2 cells into tissue resident-like cells. Our data point to the cytokine coincidentally inducing the proliferation of a small sub-population of LT-Th2 cells in which IL-33-induced transcription results in altered functional attributes associated with tissue residence, including the expression of chemokines, adhesion molecules, and genes known more widely for their expression by ILC2s. Arginine metabolism coupled to mTORC1 expression and polyamine synthesis are critical metabolic pathways underlying IL-33 induced tissue resident-like Th2 cell development.

Our data point to an important role for mTORC1 in the IL-33-dependent development of tissue resident-like Th2 cells. The role of mTORC1 as a master regulator linking metabolic cues to cell growth and activation has been extensively studied across multiple immune cell subsets (21, 37). mTORC1 activation downstream of TCR signaling is a well-established requirement for the initial activation of CD4^+^ T cells, including Th2 cells, upon antigen presentation by dendritic cells (22, 38). Our data suggest that IL-33-mediated mTORC1 upregulation represents a second wave of metabolic reprogramming necessary for Th2 cells to adapt to their tissue microenvironment. Consistent with this, mTORC1 was shown to promote the accumulation of tissue-resident memory CD8^+^ T cells in the small intestine and lungs (39, 40), highlighting the critical role of this pathway in tissue residency. It is plausible that the impaired ability to generate tissue-resident T cells following rapamycin treatment might contribute to the immunosuppressive effects of this drug to prevent organ rejection following transplants (41).

Consistent with the established role of arginine in mTORC1 activation (26) and prior findings on the use of arginine for polyamine synthesis in anti-CD3/CD28-stimulated Th2-polarized cells (32), we found that both of these arginine-dependent pathways were important in the IL-33-driven development of tissue resident-like Th2 cells, highlighting a dual role for this amino acid in promoting Th2 cell functionality. However, our results indicate that the mTORC1-dependent pathway is distinct from arginine-driven polyamine synthesis, as putrescine supplementation could not rescue the effects of arginine depletion. This indicates that both pathways independently contribute to Th2 cell functionality, and their specific contributions warrant further investigation. It is noteworthy that arginine metabolism has also been shown to play an important role in ILC2s, where the deletion of Slc7a8, a transporter also upregulated in IL-33-stimulated LT-Th2 cells, resulted in impaired mTORC1 activation, cellular proliferation and cytokine production (29). Furthermore, deletion of Arginase 1 in ILC2s dampened polyamine synthesis, proliferation and cytokine production (42).

IL-33 is emerging as a central driver of tissue residency across multiple immune cell types, including ILC2s, intestinal memory CD8 T cells, Treg cells, and macrophages (7, 8, 43, 44). Our data suggest that IL-33 plays a critical role in promoting Th2 tissue residency by inducing chemotactic and adhesion molecule expression. These findings align with studies in ILC2s (43) and adipose tissue Tregs (7, 45), where IL-33 enhances tissue localization and functionality. This highlights the conserved nature of IL-33 signaling across diverse immune cell subsets in orchestrating tissue-intrinsic immune responses.

IL-33 signals through the adaptor protein MYD88, leading to activation of the NF-κB pathway, which directly regulates gene expression, and the MAPK pathway, which stimulates AP-1 transcription factors (46). While dissecting the downstream signaling pathways of IL-33 was beyond the scope of this study, previous research has demonstrated that p38 MAPK is critical for the expression of IL-5 and IL-13 in ST2^+^ lung-resident memory Th2 cells (47). These findings suggest that the p38 MAPK pathway is likely to contribute to the enhanced cytokine secretion observed in the IL-33-stimulated Th2 cells in this study. Furthermore, this highlights the multifaceted role of IL-33 in Th2 cells, encompassing the promotion of tissue residency, cytokine secretion, and cell survival (9, 47). A better understanding of the signaling pathways downstream of IL-33 could provide further insights into the mechanisms underlying tissue-resident Th2 cell responses.

Given its pivotal role in Th2 cell function, the IL-33-arginine axis represents an attractive therapeutic target. Targeting the IL-33/ST2 axis has shown efficacy in preclinical models of allergic airway inflammation and showed promise in early clinical trials (48, 49). Our findings in Th2 cells, and findings from others working on ILC2s (29, 42), suggest that combining IL-33-targeted therapies with interventions aimed at arginine metabolism could synergistically suppress pathogenic type 2 immune responses. Such approaches may have broad applicability in treating chronic allergic diseases and other conditions driven by aberrant type 2 immunity.

## ACKNOWLEDGMENTS

This work was supported by NIH R01AI177287 (EJP), F32AI176630 (AKK), R01AI156274 (ELP), R01AI172832 (ELP), Maryland Cancer Moonshot Research Grant to the Johns Hopkins Medical Institutions (DES).

E.L.P. is a SAB member of ImmunoMet Therapeutics. E.L.P. and E.J.P are Scientific Advisors to Remedy Plan Therapeutics.

## METHODS

### Mice

C57BL/6J (the Jackson Laboratory: 000664), B6.SJL-*Ptprc^a^ Pepc^b^*/BoyJ (the Jackson Laboratory: 002014), and IL-4 reporter (4get) (18) mice were used. 4get mice were backcrossed to C57BL/6J background by Dr. Irah King at McGill University and shared with us. Mice were housed within a 12 h dark/light continuous cycle and access to water and food was *ad libitum.* All mice were maintained at the Johns Hopkins University and all corresponding animal protocols were approved by the animal care committee of the Animal Care and Use Committee (ACUC) at Johns Hopkins University. Animals used for tissue harvest or experimental procedures were aged between 7 and 12 weeks at the start of the experiment and were age- and sex-matched. Both female and male mice were used in the study.

### *Hpoly* infection

*H. polygyrus bakeri* (*Hpoly*) L3-stage larvae were prepared at the U.S. Department of Agriculture (Beltsville, USA). Mice were administered with 200 L3-stage larvae in PBS via oral gavage. For primary infection, mice were sacrificed 13 to 15 days post-infection.

### Isolation of cells from the adipose tissue

For isolation of cells from mAT, mice were euthanized and perfused with 10 ml of PBS. The adipose tissue was collected, chopped, and digested for 30 min at 37°C in fat media (DMEM, glucose [1g/L], 25 mM HEPES, 1% low fatty acid BSA, 2 mM L-glutamine, 100 U/ml P/S) supplemented with Liberase TL (Roche, 0.2 mg/ml) and DNase I (Roche, 0.25 mg/ml). After digestion, DMEM containing two mM EDTA was added and the suspension was filtered through a 70 µm strainer. Stromal vascular fraction (SVF) cells were separated from adipocytes by centrifugation and ACK lysis buffer was used to lyse red blood cells.

### T cell isolation and culture

CD4 T cells were isolated from the spleen and mesenteric lymph nodes of naïve or *Hpoly*-infected mice using MojoSort™ Mouse CD4 T Cell Isolation Kit (Biolegend, 480033). The cells were counted and plated at 0.5 million cells/ml (1.2ml in 12 well plate) in T cell media (1640 RPMI, 10% FBS, 2mM L-glutamine, 100 U/ml P/S; TCM). The cells were cultured with human IL2 (PeproTech, 200-02-1mg; 100 Units/ml) alone or together with IL-33 (R&D Systems, # 3626-ML-010/CF, 25 ng/ml) for five days. In certain experiments, alternative cytokines or cytokine combinations were used: IL-4 (PeproTech, 214-14; 10ng/ml), IL25 (R&D Systems 1399-IL-025/CF; 25ng/ml), TSLP (R&D Systems, 555-TS-010/CF; 25ng/ml). For CD3/CD28 stimulation, cells were cultured in a plate coated with an anti-CD3 antibody (5μg/ml) in TCM supplemented with anti-CD28 (BioXCell, Clone 37.51; 2 µg/ml), IL-4 (PeproTech, 214-14; 10ng/ml), IL2 (PeproTech, 200-02-1mg; 100 Units/ml), and anti-IFNy (BioXCell, clone XMG1.2; 4μg/ml). After three days, the cells were harvested, split, and cultured with IL2, IL4, and anti-IFNy. Where indicated, cells were treated with the following inhibitors: 20 nM rapamycin (Calbiochem) or 1.25mM DFMO (Enzo Life Sciences).

### ELISA

Concentrations of IL-5, IL-13, and amphiregulin in media supernatants were determined via ELISA according to the manufacturer’s protocols (R&D Systems, DY405, DY413, DY989).

### Flow cytometry

For analysis of intracellular cytokine production cells were restimulated for 4-6 hours at 37°C in TCM supplemented with 0.1ug/ml Phorbol12-myristate 13-acetate (PMA) and 1 μg/ml Ionomycin in the presence of 10 µg/ml Brefeldin A. Cells were stained in FACS buffer (PBS, 1%FCS, 2mM EDTA) with Fc block (Biolegend 101302), antibody master mix, and LIVE/DEAD Fixable Near-IR Dead Cell Stain (ThermoFisher Scientific, L34976) for 30 min at 4°C. For transcription factor analysis, the cells were fixed and permeabilized using FOXP3/transcription factor staining kit (ThermoFisher Scientific, 00-5523-00); whereas, for cytokine staining BD Cytofix/Cytoperm was used (BD Bioscience, 554722). The following fluorochrome-conjugated antibodies were used: CD45 (clone 30-F11), CD4 (clone RM4-5), TCR-β (clone H57-597), TSLPR (clone 22H9), CD44 (clone IM7), CD62L (clone MEL-14), CD198 (CCR8; clone SA214G2), CD54(ICAM; clone YN1/1.7.4), GATA3 (clone TWAJ), FOXP3 (clone FJK-16s), IL5 (TRFK5), IL13 (eBio13A), pS6 (S235/236; CST, clone D57.2.2e). Additionally, biotinylated antibodies were used for the staining ST2 (IL-33R) (MDB, clone DJ8) and amphiregulin (R&D Systems, #BAF989). Flow cytometry was performed on BD FACSymphony and analyzed using FlowJo 10.8.1.

For mitochondrial staining, cells were stained with Mitotrackers Green (50nM; ThermoFisher Scientific, M7514) and Mitotracker DeepRed (50nM; ThermoFisher Scientific, M22426) for 30min at 37C followed by staining with an antibody master mix.

For SCENITH assay, Click-iT Plus OPP Alexa Fluor 647 Protein Synthesis Assay Kit (Invitrogen) was used. Briefly, cells were incubated for 30min at 37C with oligomycin (1 µM; Sigma, 75351), 2DG (100mM; Sigma, D8375), or both inhibitors, followed by incubation with OPP reagent for another 30min. Cells were then fixed and permeabilized, and the Click-iT reagent was performed according to the manufacturer’s protocol.

In all experiments, flow cytometry was performed on BD FACSymphony and analyzed using FlowJo 10.8.1.

### Metabolomics

Polar metabolites and lipids were extracted using methanol and chloroform for two-phase extraction. The samples were spun down at full speed. The top polar layer and the bottom organic layer containing lipids were collected. The polar metabolites were incubated at −80C overnight to remove residual protein, spun down, concentrated in speed vac, and resuspended in 50% acetonitrile for analysis. Lipids were dried and then resuspended in isopronanol: acetonitrile:water (2:1:1) mixture. Metabolites were quantified by LC-MS. Chromatographic separation for polar metabolites was performed using HILIC column using polar Solvent A (20mM Ammonium carbonate, 5 uM Medronic acid in H2O) and polar solvent B: 10% polar solvent A, 90% MeCN (Acetonitrile). Chromatographic separation for lipids was performed on Zorbax Eclipse Plus C18 column using lipid solvent A (10mM Ammonium formate, 60% Acetonitrile, 40% H2O) and lipid solvent B (10mM Ammonium formate, 90% IPA, 10% Acetonitrile). For some experiments, the metabolites were extracted with acetonitrile, methanol, and water mixture (40:40:20) and submitted to General Metabolics for analysis using flow-injection mass spectrometry.

### Arginine tracing and polyamine detection

13C-Argining tracing was performed by culturing cells for five days in SILAC media supplemented with 0.2 mM L-lysine and 1.1 mM 13C-Arginine (Cambridge Isotope Lab.), 10% FCS, 2mM L-glut, 100 U/ml P/S. The metabolites were extracted using a mix of cold methanol, acetonitrile and water (50:30:20) containing 1.5% hydrochloric acid. The samples were incubated at −80°C overnight, spun down at full speed, and supernatants were collected and used for analysis on the mass spectrometer. Metabolites were quantified by LC-MS. Chromatographic separation of polyamines was performed using an Atlantis Premier BEH C18 AX Column, 1.7 µm (2.1 x 100 mm, 1.7 µm particles) using a solvent gradient of Solvent A (0.1% FA in water) to Solvent B (0.1% FA in acetonitrile).

### RNAseq

For RNAseq, 4get mice were infected with *Hpoly* and sacrificed two weeks later. CD44+GFP+ (IL4+) Th2 cells were FACS isolated from pooled spleen and lymph nodes at D0 and five days after *in vitro* culture with IL2, IL2 and IL-33, or CD3/CD28 under Th2 polarizing conditions. The cells were sorted directly into RLT buffered snap frozen. RNA extraction, library preparation using SMART-Seq® v4, and Illumina sequencing were performed at Admera Health. Sequenced libraries were processed with deepTools(Ramírez et al. 2016) v_2.0, using STAR(Kaminow et al. 2021) v_2.7.10, for trimming and mapping, and feature Counts(Liao et al. 2014) v_2.0.3 to quantify mapped reads. Raw mapped reads were processed in R (Lucent Technologies)(R Core Team 2013) with DESeq2(Love et al. 2014) v_1.36 to generate normalized read counts to visualize as heatmaps using Morpheus (Broad Institute) and determine differentially expressed genes with greater than 1.5-fold change and lower than 0.05 adjusted P-value. Gene ontology analysis was performed using DAVID Functional Annotation Bioinformatics Microarray Analysis (Jiao et al. 2012) (v_2016 and v_2021). Gene set enrichment analysis (GSEA) (50) was performed using a pre-ranked gene list generated by multiplying the sign of the fold change (negative or positive) by the −log_10_ of the adjusted *p* value. All sequencing data have been deposited in NCBI Gene Expression Omnibus (GEO) under the following accession numbers: xxxxxx

### scRNA seq data

The scRNAseq dataset was processed the same as in the original publication (9). In brief, samples were demultiplexed and aligned using Cell Ranger 2.2 (10X genomics). Read count matrices were processed, analyzed and visualized in R using Seurat v.3 (93) and Uniform Manifold Approximation and Projection (UMAP) (McInnes, L, Healy, J, UMAP: Uniform Manifold Approximation and Projection for Dimension Reduction, ArXiv e-prints 1802.03426, 2018) as a dimensionality reduction approach.

---

**Sup. Fig. 1.**
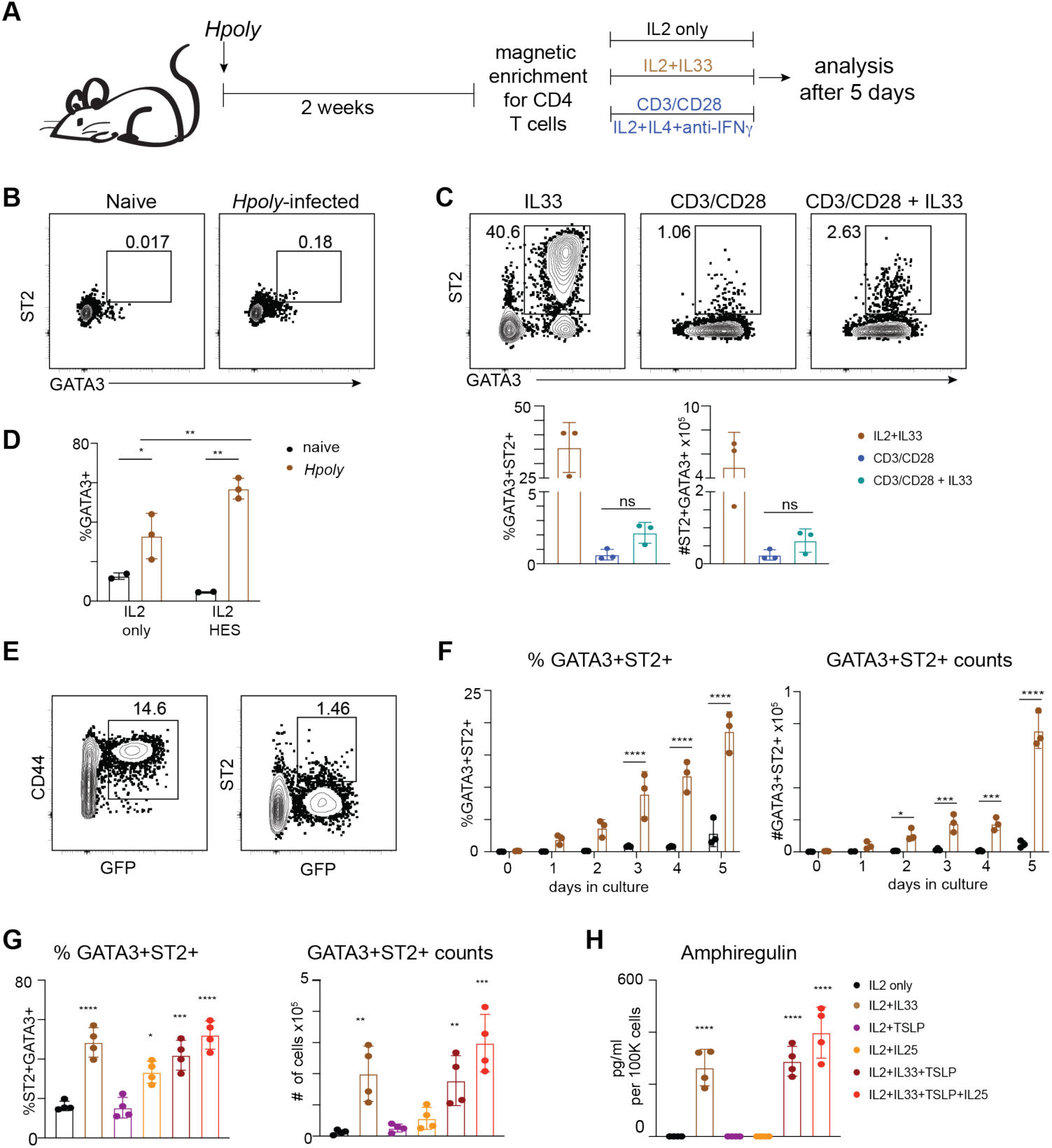
IL-33 is the primary driver of Th2_TRL_ cells. (A) A schematic of experimental design. CD4 T cells were isolated from *Hpoly*-infected mice and cultured with IL2 alone, IL2 and IL-33, or anti-CD3/CD28 under Th2 polarizing conditions for five days. (B) Representative flow plot of CD4 T cells immediately following isolation from secondary lymphoid organs of naïve or *Hpoly*-infected mice. (C) CD4 T cells from secondary lymphoid organs of *Hpoly*-infected mice were cultured under indicated conditions. Frequencies and numbers of GATA3+ST2+ cells were quantified. (D) Representative flow plot of CD4 T cells right after isolation from secondary lymphoid organs of Hpoly-infected 4get mice. (E) CD4 T cells from naïve or Hpoly-infected mice were cultured with IL2 and IL-33. (F) CD4 T cells from Hpoly-infected mice were cultured under the indicated conditions. Frequencies and numbers of GATA3+ST2+ cells were quantified after five days. (G) Abundance of amphiregulin in the culture supernatants from (F). (H) Splenocytes from naïve or Hpoly-infected mice cultured with IL2 alone or together with HES. Frequencies and numbers of GATA3+ cells was quantified. The data (B-H) are representative of at least two independent experiments. Symbols in the quantified data represent independent biological replicates. Data were analyzed by one-way ANOVA with Tukey’s post hoc test (B-G) or two-way ANOVA with Sidak post hoc test. * p ≤ 0.05, ** p ≤ 0.01, *** p ≤ 0.001.

**Sup. Fig. 2.**
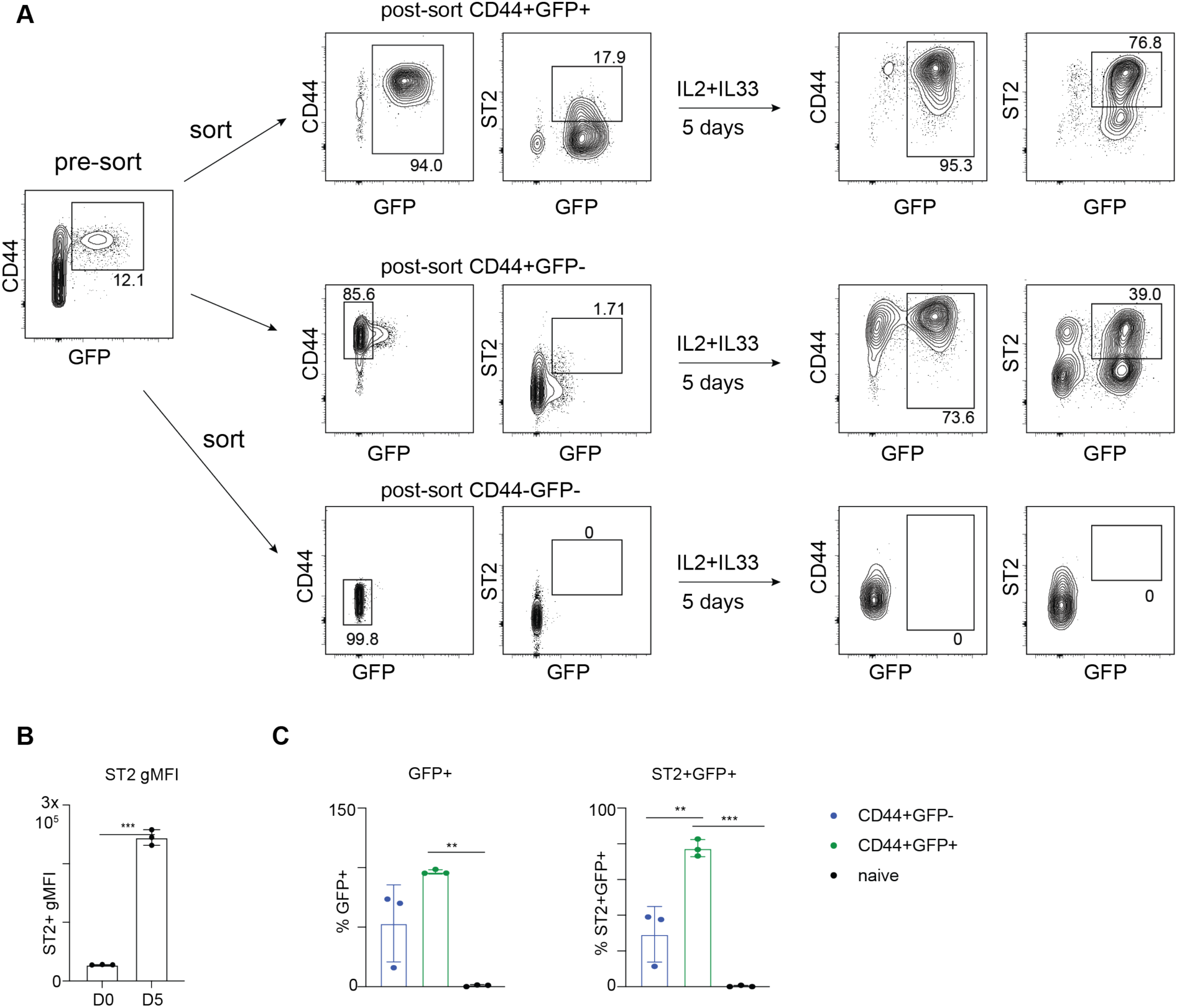
CD4 T cells from *Hpoly*-infected mice are primed to expand *in vitro*. (A) CD44+IL4+ (GFP)+ early Th2 cells, CD44+IL4-(GPF-) activated cells, and naïve CD44-GFP-cells were FACS-isolated from 4get mice infected with *Hpoly* and cultured with IL2 and IL-33 for five days. Frequencies of GFP+ and GFP+ST2+ cells (B) as well as ST2 gMFI were quantified (C). The data (A-C) are representative of at least two independent experiments. Symbols in the quantified data represent independent biological replicates. Data were analyzed by one-way ANOVA with Tukey’s post hoc test (B) or paired T-test (C). * p ≤ 0.05, ** p ≤ 0.01, *** p ≤ 0.001.

**Sup. Fig. 3.**
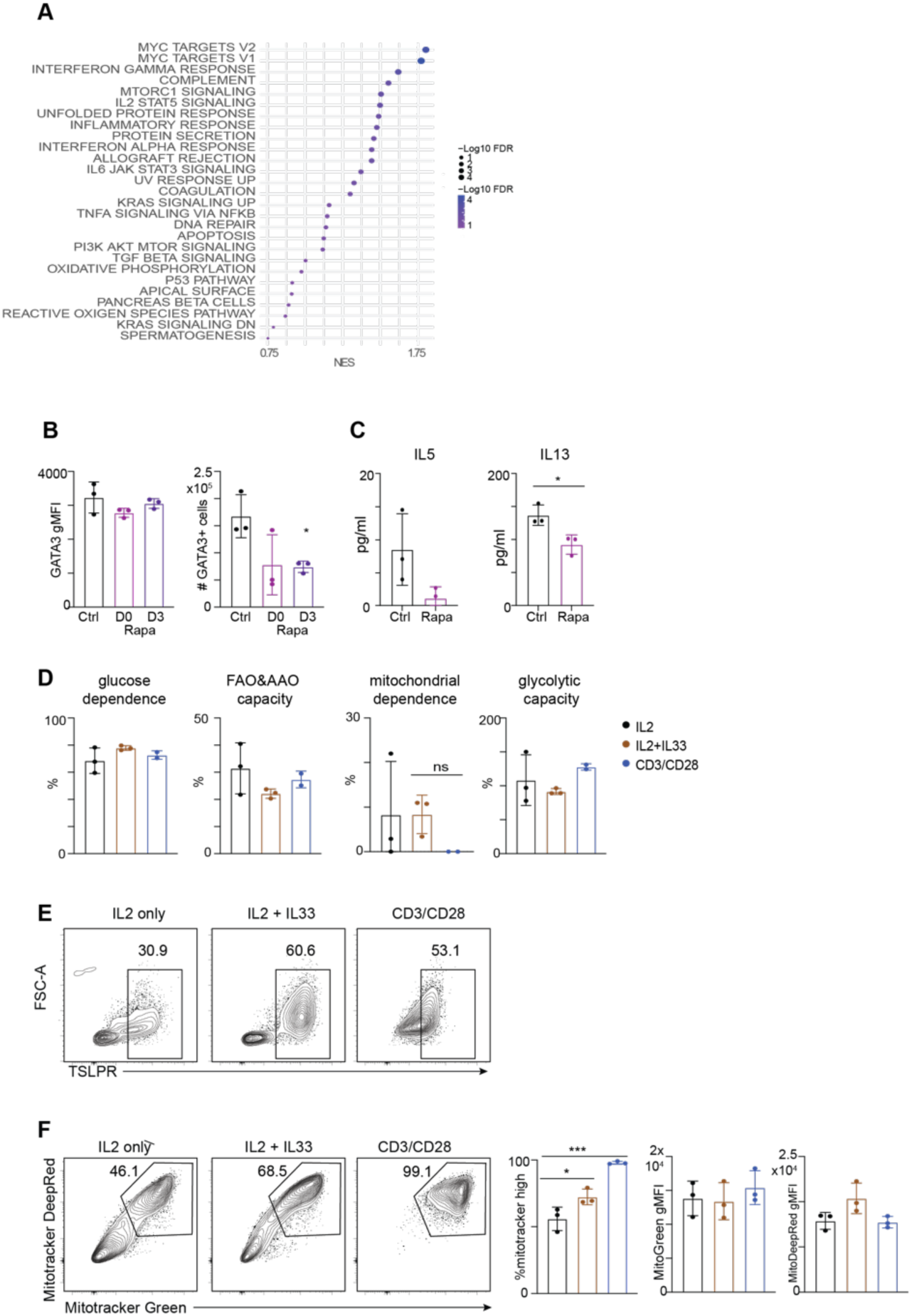
Metabolic adaptations of Th2 cells. (A) Top Hallmark pathways from GSEA for IL-33 v IL2 comparison. (B) CD4 T cells were isolated from *Hpoly*-infected mice and cultured anti-CD3/CD28 in the presence of rapamycin. The mean fluorescence intensity of GATA3 and cell numbers were quantified. (C) Levels of indicated cytokines after overnight culture of anti-CD3/CD28-stimulated cells in fresh media containing rapamycin. (D) Quantification of results from SCENITH assay in GATA3+ Th2 cells. (E) Representative flow plot showing the expression of TSLPR and ST2 after five days of in vitro culture of CD4 T cells were isolated from *Hpoly*-infected mice. (F) Representative flow plot and quantification of mitotracker green and mitotracker deep red staining after five days of culture. The data (B-F) are representative of at least two independent experiments. Symbols in the quantified data represent independent biological replicates. Data were analyzed by one-way ANOVA with Tukey’s post hoc test (B, E, F), or paired T-test (C). * p ≤ 0.05, ** p ≤ 0.01, *** p ≤ 0.001.

**Sup. Fig. 4.**
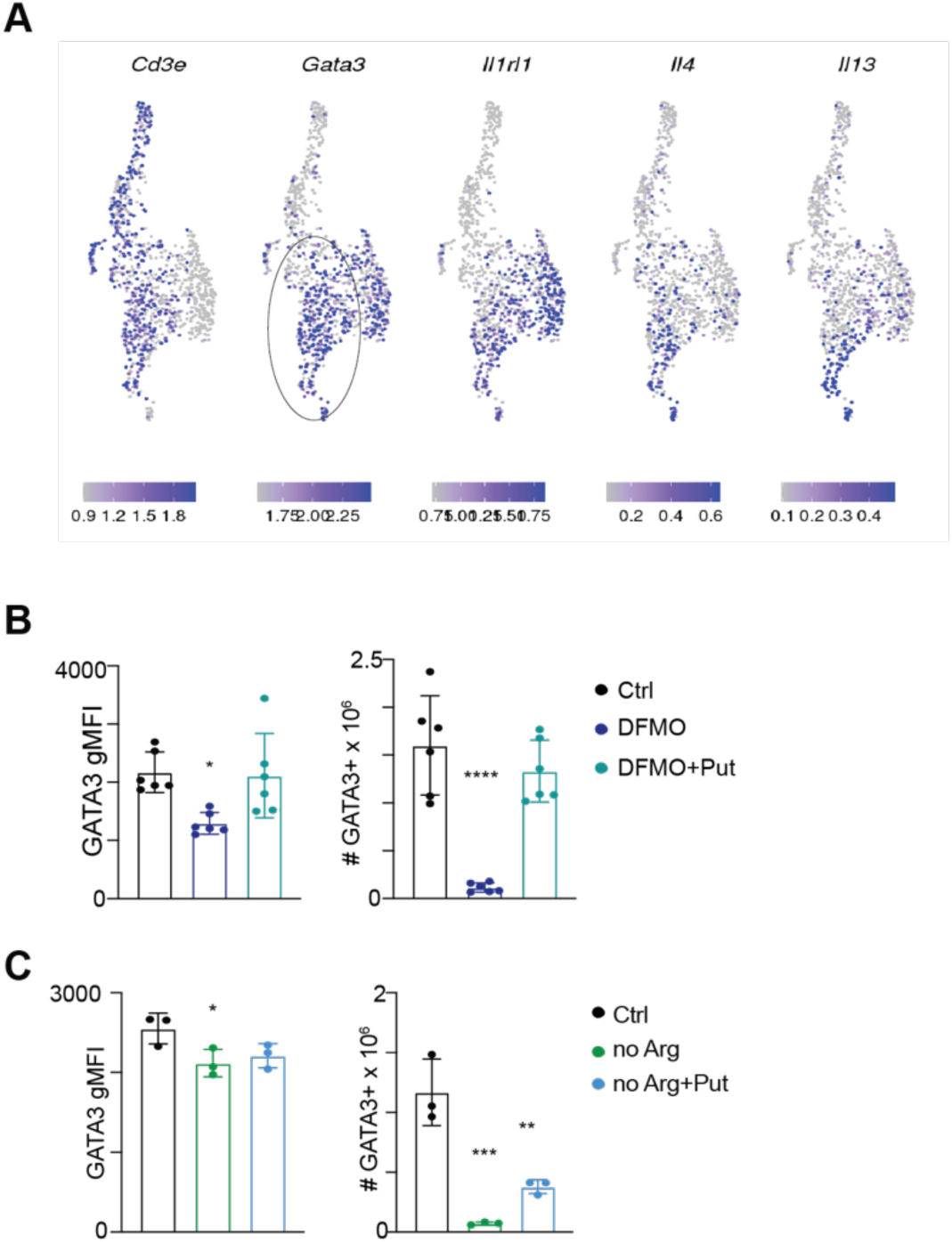
Metabolic adaptations of Th2 cells. **(A)** Expresssion of Th2 cell related genes in scRNAseq dataset of T cells isolated from mAT of *Hpoly*-infected mice identifying cluster of GATA3+ST2+ Th2 cells (circled). CD4 T cells from Hpoly-infected mice were stimulated with anti-CD3/CD28 and treated with DFMO (B) or cultured in arginine-depleted media (C). Mean fluorescence intestity of GATA3 and cells numbers were quantified. The data (B,C) are representative of at least two independent experiments. Symbols in the quantified data represent independent biological replicates. Data were analyzed by one-way ANOVA with Tukey’s post hoc test (B, C). * p ≤ 0.05, ** p ≤ 0.01, *** p ≤ 0.001.

